# Deep learning extracts MoA-specific signatures from high-throughput images of chemically and genetically perturbed *Corynebacteria*

**DOI:** 10.64898/2026.02.23.707449

**Authors:** Daniel Krentzel, Julienne Petit, Yves-Marie Boudehen, Nassim Mahtal, Elodie Sadowski, Agnès Zettor, Alexandra Aubry, Jeanne Chiaravalli, Nathalie Aulner, Stéphanie Petrella, Pedro M. Alzari, Christophe Zimmer, Anne Marie Wehenkel

**Affiliations:** Institut Pasteur, Université Paris Cité, Imaging and Modeling Unit, F-75015 Paris, France; Institut Pasteur, Université Paris Cité, CNRS UMR 3528, Bacterial Cell Cycle Mechanisms Unit, F-75015 Paris, France; Institut Pasteur, Université Paris Cité, CNRS UMR 3528, Structural Microbiology Unit, F-75015 Paris, France; Institut Pasteur, Université Paris Cité, Photonic Bio-Imaging, Centre de Ressources et Recherches Technologiques (UTechS-PBI, C2RT), F-75015 Paris, France; AP-HP, Hôpital Pitié-Salpêtrière, Service da bactériologie, Centre National de Référence des mycobactéries et de la résistance des mycobactéries aux antituberculeux, F-75013 Paris, France; Institut Pasteur, Université Paris Cité, CNRS UMR 3523, Chemogenomic and Biological Screening Core Facility, C2RT, F-75015 Paris, France; Sorbonne Université, CNRS, Inserm, Centre d’Immunologie et des Maladies Infectieuses, CIMI, F-75013 Paris, France; Machine Biophotonics Lab, Rudolf Virchow Center for Integrative and Translational Bioimaging, University of Würzburg, Germany

**Author notes:** These authors contributed equally (Daniel Krentzel, Julienne Petit).

## Abstract

Tuberculosis (TB) is the worldwide leading infectious killer due to a single pathogen and increasing antimicrobial resistance (AMR) makes it imperative to discover and develop new drugs with novel modes of action (MoAs) to treat TB infections. Phenotypic screening of chemical libraries has proven effective at identifying new compounds against bacterial pathogens. However, a major limitation of standard screens is their inability to uncover the MoA of hits thereby preventing targeted selection of compounds with novel MoAs. Linking drug perturbations to mutants from images could potentially enable to predict the targets of compounds that act through novel MoAs. Here, we develop a deep learning (DL)-based method to screen drug-treated *Corynebacterium glutamicum* (*Cglu*), a surrogate model for *Mycobacterium tuberculosis* (*Mtb*). Our DL model is based on a convolutional neural network architecture that takes high throughput images as input and is trained to distinguish between different MoAs. We show that our approach can robustly differentiate between the MoAs of established antibiotics and correctly recognise the MoA of antibiotics that were not previously seen by the DL model. We also show that inhibitors with the same and previously unseen MoA cluster together and apart from all other reference drugs, allowing for new MoA discovery. Importantly, we show that our model links images of chemical (drugs) and genetic (mutants) perturbations targeting similar pathways, thus paving the way towards mutant-based target prediction of compounds that act through novel MoAs, directly from high-content images. Finally, we explore the phenotypes induced by genetic disruption of pathways and demonstrate that features extracted with our DL model recover known biological relationships from high-throughput images alone using the cell cycle of *Cglu* as a case study, a finding with promising potential for fundamental mechanistic studies.

## Introduction

*Mycobacterium tuberculosis* (*Mtb*), a member of the suborder *Mycobacteriales* of *Actinobacteria*, is the causal agent of tuberculosis (TB). TB is amongst the top ten causes of death worldwide and the leading cause of death in patients with HIV^1^. In 2024, the World Health Organisation (WHO) revised its Bacterial Priority Pathogens List to add rifampicin-resistant *Mtb*^2^. To face an increasing burden linked to AMR, there is an urgent need for novel treatments that overcome existing resistance mechanisms in *Mtb*. Currently, TB is treated by administering a multi-drug cocktail over various months with debilitating side effects. Carrying out this drug regimen is especially challenging in low-income countries and rural settings thereby hampering the ambitious goal to eradicate TB by 2050. Reaching this goal will require a deeper understanding of the complex *Mtb* biology to enable the identification and validation of new TB targets with the aim of discovering potent *Mtb* inhibitors preferably with novel modes of action (MoAs). This could not only enable the treatment of multi and extensively drug-resistant TB but also reduce the complexity and duration of current drug regimens.

Concerningly, the development of novel antibiotics has been in steady decline over the past three decades due to low success in validating drug candidates and limited commercial attractiveness despite considerable research and development costs^3,4^. A key limitation of most drug discovery pipelines is the lack of efficiency and difficulty in reliably establishing the precise MoA of drug candidates^4–6^. Phenotypic drug discovery (PDD) is an attractive avenue to discover compounds with novel MoAs^7^. Its agnosticism to specific targets enables the discovery of previously unknown MoAs and the use of live cells removes the need for optimising chemical compounds to overcome cell permeability issues. The latter point is especially relevant for *Mycobacteriales*, since these bacteria have a unique and complex multi-layered cell wall composed of three distinct macromolecules: peptidoglycan (PG), arabinogalactan (AG) and mycolic acids (MA)^8^. These macromolecules form a formidable permeability barrier to most chemical compounds including potential antimicrobials^9^.

In recent years, developments in image-based assays (e.g. Cell Painting^10^) and image analysis methods (e.g. CellProfiler^11^) that quantitatively assess cellular phenotypic changes induced by molecules have led to an explosion in image-based PDD screens^12^. Instead of relying on mono-parametric readouts like optical density (OD), these methods enable a holistic assessment of both cellular and sub-cellular changes by extracting hand-crafted features from segmented cells. This allows the identification of compounds with subtle effects which can then be further optimised by medicinal chemists. More importantly, these image-based screens address a key limitation of growth inhibition assays, namely MoA recognition. Successful MoA recognition of molecules from images alone has been shown in various eukaryotic screens and pioneering work by Nonejuie *et al.* and Zoffmann *et al.* has demonstrated that such methods can also be applied to bacterial screens^13,14^. For instance, in a recent image-based phenotypic screen of *Acinetobacter baumannii*, such an approach was crucial to identify a novel class of antibiotics targeting the lipopolysaccharide transport machinery^15^.

Directly applying these approaches to screen for TB drugs is however challenging, since *Mtb* exhibits a high degree of cell-to-cell variability even in the absence of drug exposure. Prior work has thus sought to account for the heterogeneous response of individual *Mtb* cells either by specifically defining hand-crafted features that account for this cell-to-cell variability^16^ or by training a type of deep learning (DL) model called a convolutional neural network (CNN) to predict drug MoAs from images of drug-treated *Mtb* without cell segmentation^17^. While these methods enable MoA recognition of hit compounds with respect to known compounds, they are unable to identify drug targets of active compounds with novel MoAs. A promising avenue to overcome this limitation is to assume that drug treatments and genetic mutations result in similar phenotypes if they affect the same proteins or pathways. Along these lines, prior work has shown that hand-crafted features obtained from images of CRISPRi *Mycobacterium smegmatis* (a non-pathogenic surrogate model of *Mtb*) mutants and drug-treated bacteria tend to cluster according to the targeted pathways^18^.

Moreover, using non-pathogenic surrogate models of *Mtb,* like *M. smegmatis,* to study the effect of TB drugs can drastically speed up and simplify experimental protocols, since *Mtb* is highly pathogenic which requires experiments to be carried out within high biosafety level (BSL-3) labs, and slow-growing (24h doubling time) thus limiting throughput. However, the genome of *M. smegmatis* is almost two times larger than that of *Mtb* (7 Mbp and 4.4 Mbp respectively) with larger functional redundancy, making the study of specific cellular pathways more challenging. We therefore use another non-pathogenic surrogate model for *Mtb* called *Corynebacterium glutamicum* (*Cglu*), a member of the *Mycobacteriales,* with a short doubling time of about 1h and a genome size comparable to that of *Mtb* (3Mbp). Moreover, *Cglu* and *Mtb* share common core mechanisms related to cell division and cell wall synthesis, as well as a complex multi-layered cell wall^19,20^. A key advantage of *Cglu* is its ability to withstand the loss of its outer cell envelope layers thereby enabling the study of its biosynthetic proteins^21^,which are potential drug targets. Most importantly, for the context of a TB drug screen, *Cglu* has been shown to respond to antibiotics and in particular TB drugs that target core conserved mechanisms^22,23^.

Here, we present an image-based high-throughput TB-drug discovery pipeline that links drug-treated and mutated *Cglu* strains where the same or a similar pathway has been disrupted. To that end, we acquire high-throughput images of *Cglu* exposed to several known TB drugs and use these images to train a custom CNN for the task of predicting antibiotic MoAs from images alone. We show that our model successfully predicts targets of non-canonical TB drugs and detects and distinguishes compounds with previously unseen MoAs. We furthermore demonstrate that our DL model can extract meaningful phenotypic signatures from images of drug-treated bacteria and mutants in which the same pathway has been disrupted. This allows for the association of genetically and chemically induced phenotypic changes directly from images paving the way towards mutant-based target deconvolution. Finally, we show that our DL model provides insights into the function of genes affecting the cell cycle by clustering images of mutants belonging to similar pathways. Taken together, our approach presents a promising avenue for image-based phenotypic screening in bacteria, mutant-based target deconvolution of novel compounds and investigating the role of bacterial genes directly from images.

## Results

### Bacterial high-throughput image acquisition

To obtain a dataset for training and testing our DL model, we acquired high-throughput images of a *Cglu* strain (ATCC13032) grown in controlled medium and exposed bacteria to 18 established drugs with five different modes of action (MoAs) associated with seven different biological pathways (**Table 1**). Drugs were grouped according to modes rather than mechanisms of action, since not all drugs have a clearly identified and/or unique protein target) outlined in. To train our DL model, we selected a subset of these antibiotics belonging to five different MoAs: (i) ethambutol inhibits the arabinosyltransferases involved in the synthesis of the AG layer, a critical component of the mycobacterial cell envelope^24,25^; (ii) amoxicillin, carbenicillin and cefotaxime are beta-lactam compounds that target penicillin-binding proteins (PBPs) involved in PG synthesis^26^; (iii) the two fluoroquinolones (FQs) ciprofloxacin and moxifloxacin, and the aminocoumarin novobiocin inhibit DNA gyrase – the sole topoisomerase in *Cglu* responsible for regulating DNA supercoiling and decatenating daughter chromosomes^27^; (iv) clarithromycin, doxycycline and linezolid target the ribosome^28^ and (v) clofazimine which is thought to target the respiratory chain and has also been shown to affect membrane homeostasis in *Mtb*^29^ and *Staphylococcus aureus*^30^.

**Table 1:** Modes of action of antibiotics from training and testing dataset.

| Mode of action | Antibiotic | In training data? | MIC ($\mu\text{M}$) | References |
| --- | --- | --- | --- | --- |
| Cell wall (PBP) | Amoxicillin | ✓ | 0.342 | Hennart *et al.*<sup>51</sup> |
|  | Carbenicillin | ✓ | 0.661 | This work |
|  | Cefotaxim | ✓ | 0.263 | Hennart *et al.*<sup>20</sup> |
|  | Ampicillin |  | 0.358 | This work |
| Cell wall (AG) | Ethambutol | ✓ | 4.895 | This work |
| DNA Gyrase | Ciprofloxacin | ✓ | 0.755 | Gedeon *et al.*<sup>22</sup> |
|  | Moxifloxacin | ✓ | 0.299 | Wormser *et al. in prep* |
|  | Novobiocin | ✓ | 26 | Wormser *et al. in prep* |
|  | Gepotidacin |  | 0.446 | Gedeon *et al.*<sup>22</sup> |
|  | BDM71403 |  | 0.268 | Gedeon *et al.*<sup>22</sup> |
| Ribosome | Clarithromycin | ✓ | 1.337 | This work |
|  | Doxycycline | ✓ | 0.270 | This work |
|  | Linezolid | ✓ | 2.964 | This work |
| Cell Membrane | Clofazimine | ✓ | 2.112 | This work |
| RNA polymerase | Rifampicin |  | 0.146 | This work |
|  | Rifabutin |  | 0.295 | This work |
| Folate synthesis | Trimethoprim |  | >64 | This work |
|  | Sulfamethoxazole |  | >64 | This work |

For each of the above-mentioned drugs, we determined or used published minimum inhibitory concentrations (MICs) (**Table 1; Materials and Methods**) and exposed bacteria to four different concentrations (0.5, 1, 5 and 10xMIC). Drugs were distributed in a random layout onto six 96-well plates with an acoustic liquid handler (Beckman Coulter Echo 550) to decorrelate conditions from well-plate position (**Supp. Fig. 1**). We exposed bacteria to drugs for 16h starting in early exponential phase. Bacteria were stained for membrane (FM4-64) and nucleoid (Hoechst) and fixed with ethanol. To ensure consistent densities between treatment conditions, OD measurements were used to normalise the number of bacteria in each well prior to transferring them to imaging plates. Then, 50 three-channel (brightfield, FM4-64 and Hoechst) images per condition (i.e. combination of concentration and antibiotic) were acquired on a high-content screening system (Revvity Opera Phenix Plus) in widefield mode with a 63x water-immersion objective. In total, six biological replicates were acquired on six separate plates (**Fig. 1a**).

**Fig. 1.**
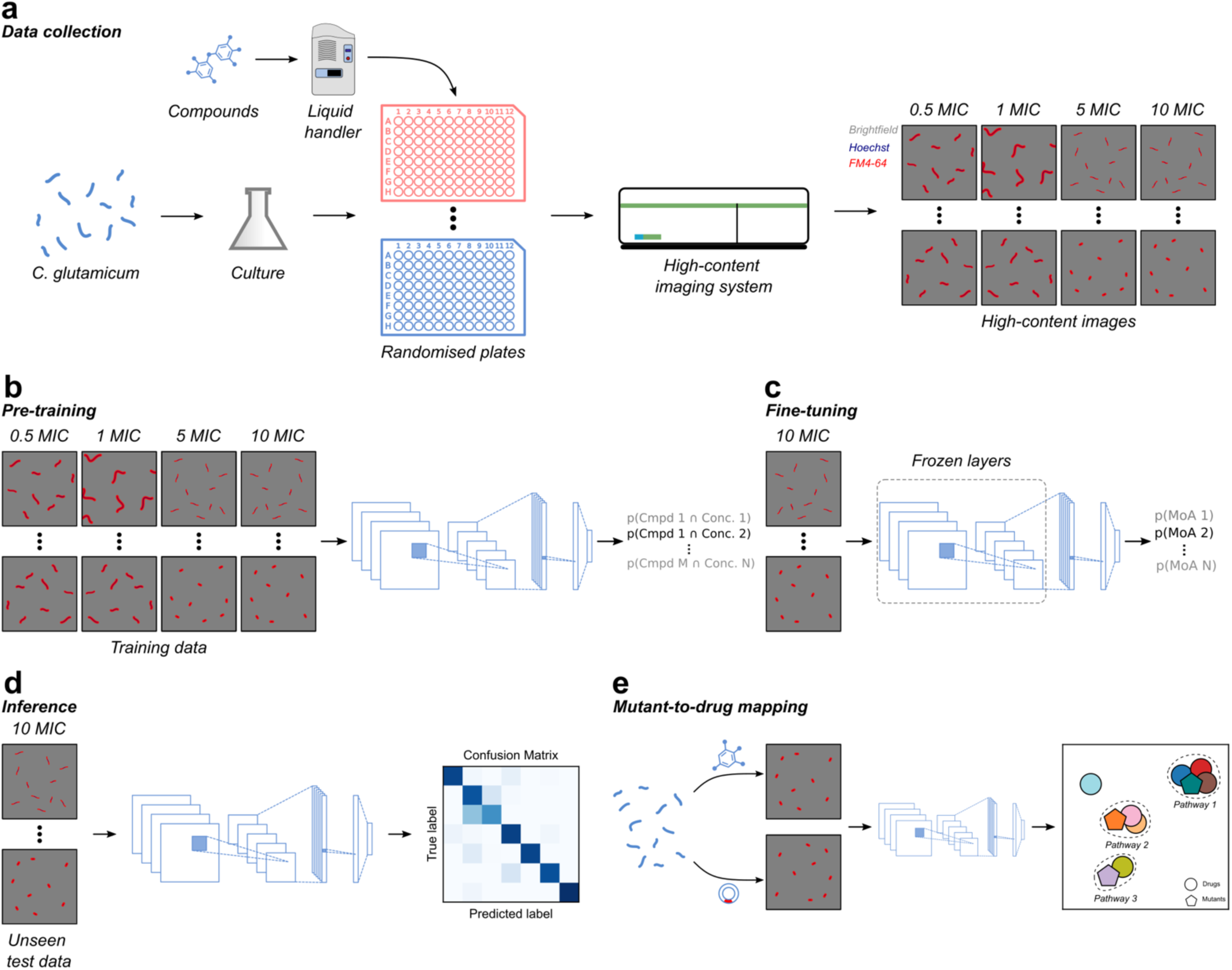
Dataset acquisition and model training. (**a**) Antibiotic compounds were distributed with randomised plate layouts at four different concentrations (0.5xMIC, 1xMIC, 5xMIC and 10xMIC) onto six 96-well plates with an acoustic liquid handler. *Cglu* cultures were grown for 16 h in six biological replicates and incubated with antibiotics on the previously prepared 96-well plates. Bacteria were then fixed with ethanol, stained with FM4-64 (cell membrane) and Hoechst (DNA), and imaged on a high-content microscope with a 63x water-immersion objective in three channels (brightfield, FM4-64 and Hoechst). (**b**) high-throughput images across all concentrations from five out of six plates were used to pre-train a custom DL model to predict the combination of antibiotic and concentration from images alone. (**c**) After pre-training, convolutional layers were frozen, and images of bacteria exposed to antibiotics at 10xMIC were used to fine-tune the fully connected layers. During fine-tuning, we used a triplet loss to explicitly promote the formation of clusters by MoA in latent space, a centre loss to increase tightness of clusters and a weighted cross-entropy loss for prediction. (**d**) Performance quantification and further analysis was conducted on unseen images from a hold-out test plate to ensure robustness to biological and technical variation. (**e**) Images of mutants and drugs were then jointly embedded using the previously trained DL model to link chemical and genetic perturbations by pathway.

### Model architecture and training strategy

To obtain high-dimensional feature vectors from bacterial high-throughput images in an end-to-end fashion, we modified a DL model that we previously developed to recognise the MoA of antibiotics from images of *E. coli*^31^. First, we pre-trained our model using five out of six plates (biological replicates) to predict combinations of drugs and concentrations directly from images as described in our previous work^31^ (**Fig. 1b; Materials and Methods**). Then, we added a fine-tuning step where the convolutional layers were frozen and only parameters in the fully connected layers of our model were updated (**Fig. 1c**). For fine-tuning, we directly trained our model to predict drug MoAs using images of bacteria exposed to antibiotics at the highest concentration (10xMIC) to increase the likelihood of bacteria showing discriminative phenotypes. We found that the addition of this fine-tuning step during training improved clustering by MoA (**Supp. Fig. 2**). We then tested the performance of our DL model on a sixth hold-out test plate (biological replicate) with a different randomised plate layout (**Supp. Fig. 1**) to ensure robustness to positional effects and biological variation. To compare phenotypes between different perturbations, we obtained high-dimensional feature vectors from the penultimate layer of our model (**Fig. 1d**).

**Fig. 2.**
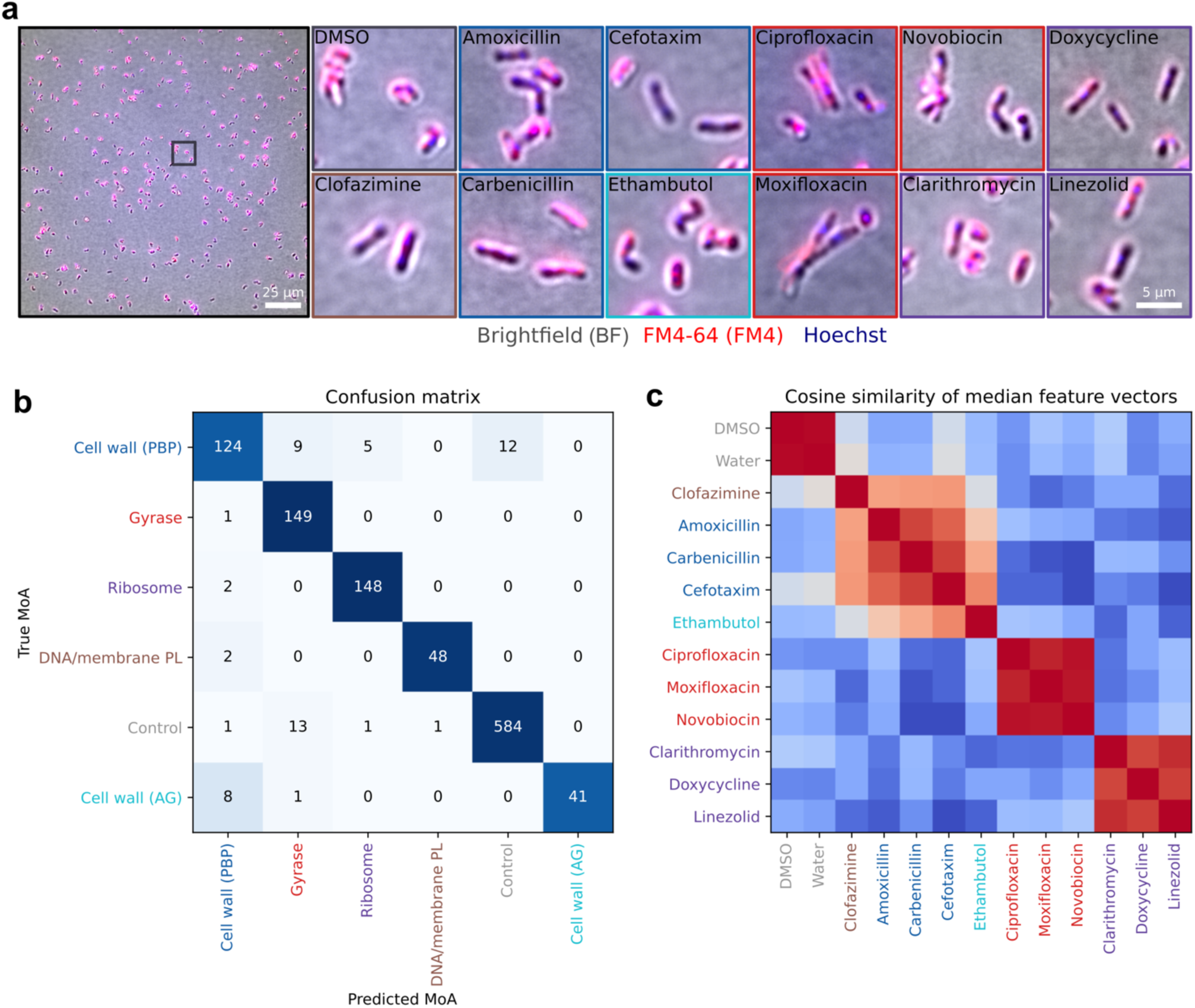
Deep learning recognises MoAs of antibiotics from images alone. (**a**) Representative example images of *Cglu* bacteria exposed to antibiotics at 10xMIC and acquired on a high-content microscope with a 63x water-immersion objective are shown with scale bars. Brightfield is shown in greyscale, cell membrane (FM4-64) in red and DNA (Hoechst) in blue. (**b**) Confusion matrix obtained from a DL model trained with three-channel images (brightfield, FM4-64 and Hoechst) of bacteria exposed to antibiotics at 10xMIC and tested on unseen images from a hold-out test plate. (**c**) Cosine similarity computed on median feature vectors obtained from images of drug-treated bacteria shows high similarity (in red) of antibiotics by MoA.

To explore if we could directly link chemical to genetic perturbations from images, we prepared a second dataset based on biological triplicates of plates containing seven mutant strains and *Cglu* bacteria exposed to seven different antibiotics (**Table 2**). After acquiring high-throughput images as described above, we used the DL model, that we previously trained to predict the MoA of antibiotics outlined in **Table 1**, to obtain feature vectors for all conditions which we used to directly link drug perturbations to bacterial mutant strains (**Fig. 1e**).

**Table 2:** Description of dataset shown in Fig. 4 of disrupted pathways of mutants and drugs. Full mutant description can be found in Supp. Table 1.

| Mode of Action | Mutant name | Drugs |
| --- | --- | --- |
| DNA segregation | *gyrA*<br>*gyrB* | BDM71403<br>Ciprofloxacin<br>Gepotidacin<br>Moxifloxacin |
| Cell wall/division defect | *sepF*<br>$\Delta glp$<br>$\Delta glpR$<br>$\Delta steAB$ | Ampicillin<br>Carbenicillin |
| Cell wall/elongation defect | *wag31* | Ethambutol |

### Deep learning identifies the MoA of previously seen drugs with high accuracy directly from images

We first validated that our DL model could distinguish the MoA of established drugs. To that end, we used high-throughput images of *Cglu* bacteria exposed to eleven antibiotics, the majority of which are used in clinics to treat TB (**Table 1**). We used the DL model to predict the MoA from images of bacteria exposed to antibiotics at 10xMIC on six rotations of the training and testing plates on the subset of the eleven antibiotics that were included in the training set. Our model achieved a per-FoV (field of view) classification accuracy of 86.78±7.90% cross-validated across six hold-out test replicates on these previously seen drugs. To verify that the DL model learned to extract more similar feature vectors for drugs belonging to the same MoA, we computed a cosine similarity matrix using the median feature vectors obtained for each drug treatment (**Fig. 2c**). As expected, we found that compounds with similar MoAs exhibited high cosine similarity. Interestingly, despite training the DL model to consider clofazimine and ethambutol as distinct classes, median feature vectors for both conditions exhibited similarity to drugs targeting PBPs. Indeed, cells treated with clofazimine exhibit an elongated cell length similar to beta-lactam inhibition (**Supp. Fig. 3**). Plotting feature vectors with UMAP revealed a broad cluster encompassing drugs targeting the cell envelope in which ethambutol and clofazimine can be discriminated from PBP inhibitors (**Supp. Fig. 2c, d**). This indicates that our DL model learned to pick up common phenotypic features related to cell envelope perturbations while also encoding discriminative features associated with individual MoAs, namely disruption of PG, AG and membrane homeostasis.

**Fig. 3.**
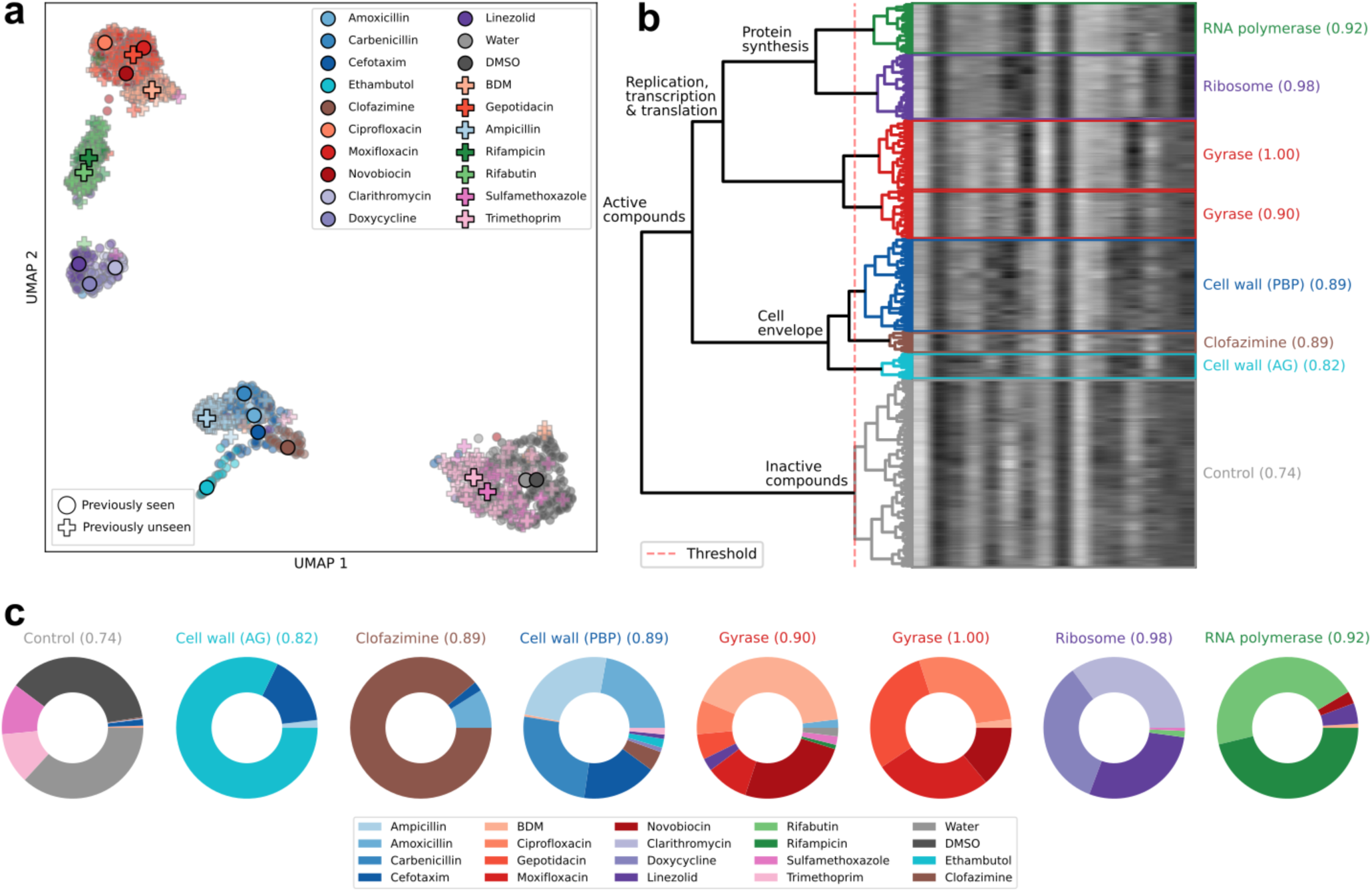
Deep learning correctly identifies drug targets of previously unseen compounds. (**a**) UMAP of feature vectors with previously seen drugs shown as dots and previously unseen drugs with crosses. Large dots/crosses correspond to median feature vectors. Four of the previously unseen drugs belong to two previously unseen MoAs inhibiting folic acid synthesis and RNA polymerase. Sulfamethoxazole and trimethoprim (folic acid synthesis) were included as negative controls, since these drugs need to be administered in combination for effects to be observed. (**b**) Hierarchical clustering on feature vectors with the threshold set such that negative controls (Water and DMSO) cluster. Clusters are annotated by the most common MoA with the proportion of drugs belonging to a given MoA shown in brackets. (**c**) Pie-charts showing the composition of each cluster by drug with colours corresponding to the drug identity.

Finally, we investigated the robustness of the DL model to changes in concentration using images of bacteria exposed to 3 concentrations below 10xMIC (0.5, 1 and 5xMIC; **Supp. Fig. 4a**). At low concentrations (0.5 and 1xMIC), most previously seen drugs were classified as control except for three PBP inhibitors (ampicillin, carbenicillin and cefotaxime) and one DNA gyrase inhibitor (novobiocin), which were correctly classified regardless of concentration, as they exhibited strong MoA-specific phenotypic changes even at low concentrations. At a concentration of 5xMIC, almost all drugs with a previously seen MoA were correctly classified suggesting that drug phenotypes stabilised. The only exception was clofazimine which was classified as a PBP inhibitor at 5xMIC (in line with the high cosine similarity observed between clofazimine and PBP inhibitors at higher concentrations in **Fig. 2c**). Interestingly, we also observed that compounds belonging to the same MoA followed similar dose-dependent trajectories in feature space (**Supp. Fig. 4b,c**).

**Fig. 4.**
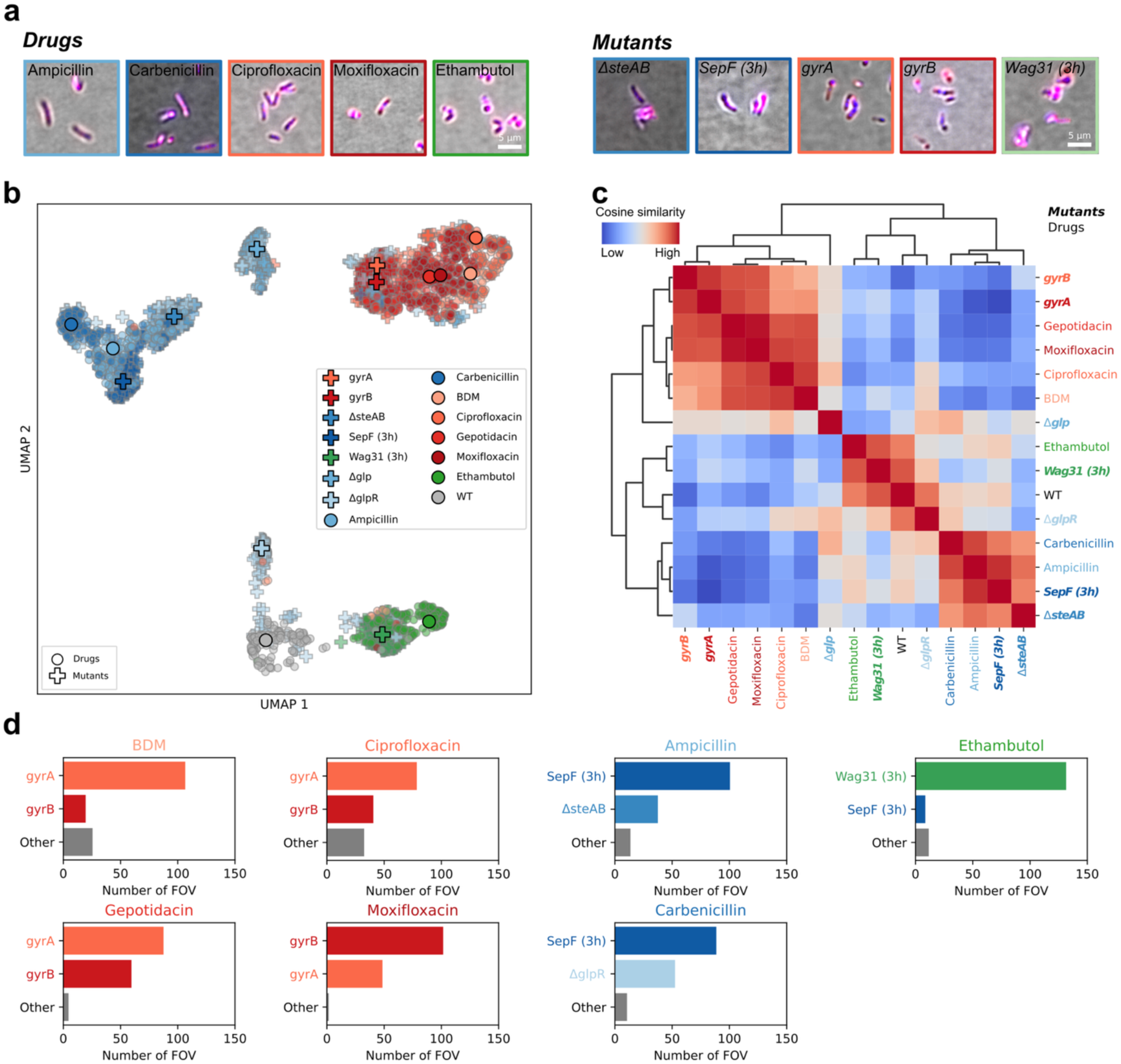
Deep learning links drugs to mutants from images alone. (**a**) Image crops of drug-treated and mutant bacteria with scale bars. (**b**) UMAP of feature vectors coloured by condition with small dots corresponding to images of bacteria treated with antibiotics at 10xMIC and crosses to mutants. Large dots/crosses correspond to median feature vectors computed across three technical replicates. (**c**) Cosine similarity matrix of median feature vectors with hierarchical clustering shows mutants and bacteria treated with antibiotics disrupting similar pathways clustering. (**d**) A MLP was trained to predict affected pathways from feature vectors of mutants. Predictions on images of drug-treated bacteria were then obtained across three replicates.

Taken together, these results show that our DL model can robustly classify drugs by their MoA from images alone and recognises similarities between drugs targeting the same or related pathways. Moreover, MoA predictions above 5xMIC are robust which indicates that accurate MoA determination of putative hit compounds could be feasible at a single concentration directly from high-throughput images.

### Deep learning identifies MoAs of previously unseen drugs and detects novel MoAs

Having verified the ability of our DL model to correctly recognise the MoA of drugs seen during training, we wanted to check its performance on unseen candidate drugs using the previously seen antibiotics as reference compounds (**Table 1**). We picked candidate drugs from three different categories: (i) compounds which exhibit no detectable activity, (ii) active compounds where the MoA, but not the drug, was present in the training data and (iii) active compounds with a previously unseen MoA. For the first category, we selected trimethoprim and sulfamethoxazole – two folic acid synthesis inhibitors that have bacteriostatic effects and are typically administered in combination^32,33^ and are inactive in *Cglu* at the concentrations that we tested (**Supp. Fig. 5**). For the second category, we used three compounds that are relevant for *Mycobacteriales* species but not typically used to treat TB infections in clinics: two novel bacterial topoisomerase inhibitors (NBTIs) targeting the DNA gyrase (gepotidacin and BDM71403) and one drug targeting PBPs (ampicillin). NBTIs differ from FQs in their mechanism of action through a distinct topoisomerases binding mode: while two FQ molecules intercalate into the cleaved DNA and interact with the GyrB and GyrA subunits to block the DNA religation step, a single NBTI molecule intercalates into the single-stranded cleaved DNA and binds only to GyrA, trapping the gyrase in a DNA-bound conformation^27^. For the third category, we chose rifampicin and rifabutin which target the RNA polymerase.

**Fig. 5.**
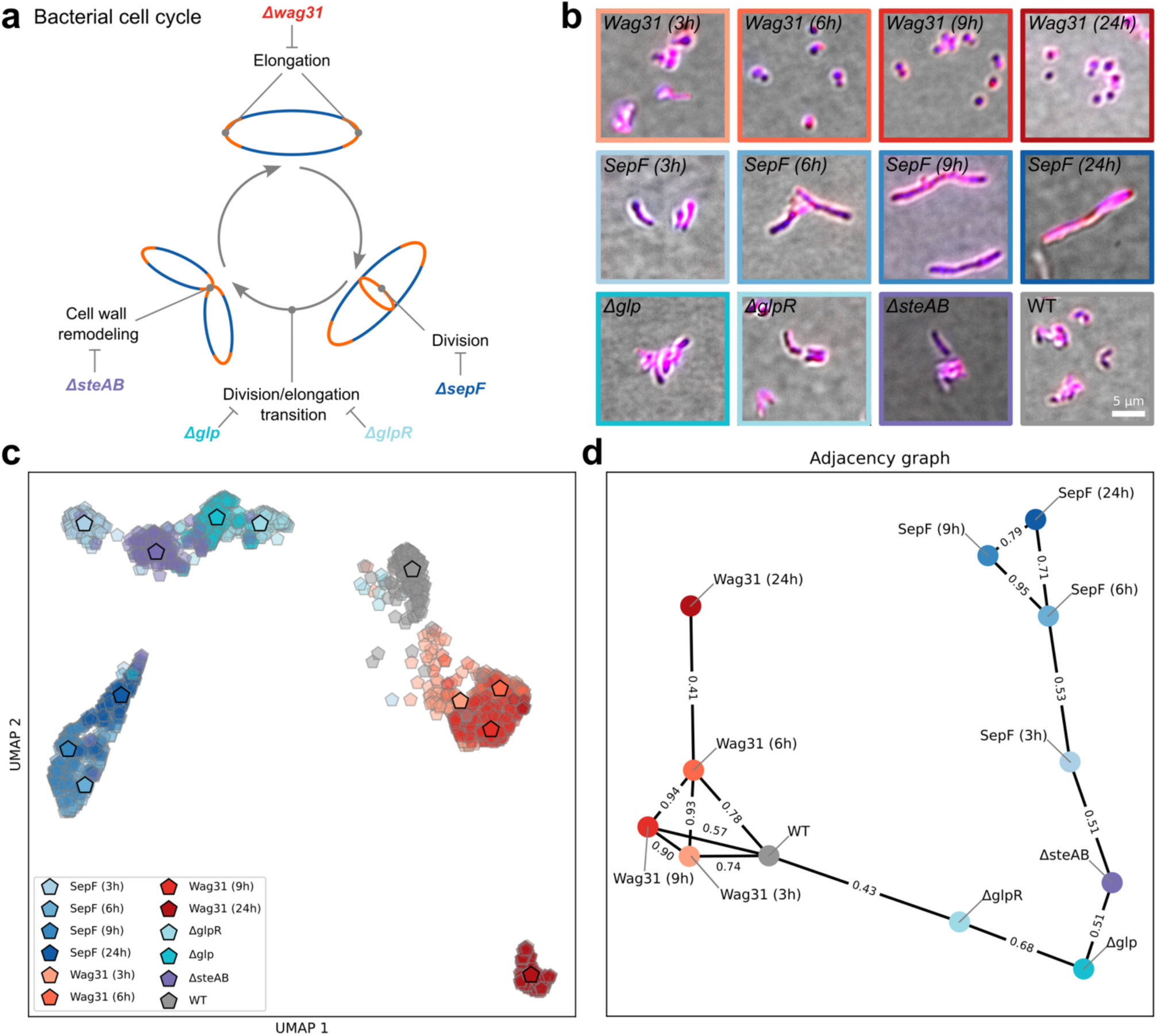
Images of bacterial cell cycle mutants cluster by disrupted pathway. (**a**) Diagram showing the bacterial cell cycle and pathways affected by mutants. (**b**) Representative image crops shown for each mutant with scale bars. (**c**) UMAP of image embeddings coloured by mutants computed on three technical replicates with small pentagons corresponding to individual images and large pentagons to median feature vectors. (**d**) Adjacency graph computed from a cosine similarity matrix (**Supp. Fig. 7c**) computed on median feature vectors and coloured by mutants showing connections between mutants with the most similar feature vectors. Values on edges correspond to cosine similarities between mutants. The threshold for the adjacency matrix was set such that each node was connected by at least one edge.

First, we used our DL model to obtain feature vectors of these previously unseen candidate drugs at 10xMIC and jointly visualised them on a UMAP plot with feature vectors of reference drugs that were previously seen during training (**Fig. 3a**). We found that candidate drugs with a previously seen MoA clustered tightly with reference drugs belonging to the same MoA. For instance, gepotidacin and BDM71403 correctly clustered with DNA gyrase inhibitors, while ampicillin clustered with the other beta-lactams. The two inactive drugs trimethoprim and sulfamethoxazole co-localised with negative controls, while rifampicin and rifabutin – the two drugs with a previously unseen MoA – formed a separate cluster.

To further quantify these observations, we performed hierarchical clustering (**Materials and Methods**) on feature vectors (**Fig. 3b, c**). The negative control feature vectors (Water and DMSO) were used to define the clustering threshold. We observed two major clusters consisting of inactive and active compounds. The latter was in turn composed of two groups, one encompassing drugs disrupting replication, transcription and translation, and the other including drugs targeting the cell envelope, in agreement with the UMAP representation shown in **Fig. 3a**. The replication, transcription and translation cluster was further split into compounds targeting protein synthesis and DNA gyrase. This indicates that the alterations in bacterial phenotypes resulting from drug exposure indeed mirror functional similarities in the pathways they affect.

Then, we used our DL model to obtain MoA predictions from candidate drugs (**Supp. Fig. 6a**). Images of candidate drugs where the MoA was included in the training data were all correctly classified. Sulfamethoxazole and trimethoprim were mostly classified as control, in line with these drugs being individually inactive at the concentrations used (**Supp. Fig. 5**). The two drugs with a previously unseen MoA (rifampicin and rifabutin) were split between two distinct “known” MoA classes: DNA gyrase and ribosome inhibition, consistent with our finding in **Fig. 3b** that our DL model clusters drugs targeting these related pathways. Next, we computed outlier scores on feature vectors of candidate drugs using a local outlier factor algorithm, an approach that we developed in prior work for detecting drugs with novel MoAs in *E. coli*^31^ (**Materials and Methods**). As expected, both rifampicin and rifabutin were characterised by significantly higher outlier scores than drugs with a previously seen MoA (**Supp. Fig. 6b**).

Overall, these results show that our DL model generalises to previously unseen drugs and correctly matches active drugs to their correct MoA when other drugs with the same MoA are present in the training data and classifies inactive drugs as control. Importantly, images of bacteria exposed to drugs with the unseen MoA (RNA polymerase inhibitors) cluster together and apart from all other conditions thus paving the way for automated detection of novel classes of inhibitors directly from images. Our results also show that phenotypes of compounds targeting similar pathways can be grouped into functionally meaningful clusters from images alone.

### Linking drugs to mutants from high-throughput images

If a drug acts on a single gene product or biological pathway, a mutant strain genetically disrupted for the same gene or a gene involved in the same pathway should lead to a phenotype recognised as similar by our DL model. Based on this hypothesis, we asked whether we could identify the target of compounds by comparing drug-induced phenotypes to phenotypes of mutants of the genes encoding these targets. As a proof of concept, we chose mutants affecting four major biological pathways of the cell cycle of the polar growing *Cglu* (**Table 2; Supp. Table 1**): (i) strains enabling inducible repression of the genes *gyrA* and *gyrB*, which respectively encode the two subunits GyrA and GyrB of the essential DNA gyrase complex^34^ that modifies DNA topology and is crucial for DNA replication and segregation, (ii) a strain for inducible repression of *sepF,* the gene coding for the membrane anchor of the tubulin homologue FtsZ^35^ that is essential for Z-ring assembly and required for septum formation and cell wall biosynthesis, (iii) three different deletion mutants (∆*steAB*^36^,∆*glp*^37^ and ∆*glpR*^37^) of genes involved in the divisome-elongasome transition at the septum, and (iv) a conditional mutant *wag31*^38^ encoding for the essential polar elongasome scaffold Wag31, which leads to disrupted polar cell wall biosynthesis. In addition to these four groups of mutants, we also included *Cglu* bacteria exposed to seven drugs at 10xMIC which target similar pathways (**Table 2**). We then prepared three plates with randomised layouts (**Materials and Methods**) containing the above-described mutants and drug-treated bacteria and imaged them as previously detailed (**Fig. 4a**).

First, we extracted feature vectors from the images of mutants and drug-treated bacteria in this new data set, using our previous DL model without retraining (**Table 1**) and plotted them with UMAP (**Fig. 4b**). Feature vectors obtained from the three replicates were in good agreement for all conditions showing that our DL model was robust to changes in well positions and plate-to-plate variations (**Supp. Fig. 7a**). We additionally computed a cosine similarity matrix on median feature vectors for all conditions and performed hierarchical clustering (**Fig 4c**; **Materials and Methods**). Our analysis revealed three mixed clusters of drugs and mutants disrupting similar pathways. First, the four anti-gyrase drugs (BDM71403, ciprofloxacin, gepotidacin and moxifloxacin) and the two DNA gyrase mutants (*gyrA* and *gyrB*) formed a tight cluster (**Fig. 4b,c**). This is expected, since these drugs directly target the protein underlying the gyrase-depletion mutant. More surprising was the fact that drugs interfering with cell wall biosynthesis (beta-lactam PBP inhibitors, ampicillin and carbenicillin) clustered with two cell-division mutants (*sepF*, *∆steAB*). While the proteins produced by these genes are not the direct drug targets of these drugs, they are nonetheless directly linked to PG biosynthesis. For instance, ampicillin and carbenicillin clustered with an early time-point *sepF* depletion mutant, as well as the *∆steAB* deletion strain. SepF is the membrane anchor of the tubulin-homologue FtsZ and thus essential for septum formation through a ring-like structure (Z-ring) that positions the PG biosynthetic machinery at the division site^35^, while the SteAB complex regulates RipA, the major endopeptidase responsible for PG remodelling at cytokinesis^36^. Hence, these two mutants, although not being depleted for PBPs (the direct targets of the ampicillin and carbenicillin), are involved in regulating PG synthesis at the septum and form a physiologically relevant cluster with beta-lactam antibiotics. The AG inhibitor ethambutol, known to disrupt the ability of cells to elongate from their poles^23,39^, consistently clustered with the *wag31* mutant which lacks the elongasome scaffold Wag31, essential for rod shape and pole maintenance.

The relative consistency between feature vectors of mutants and drugs affecting related pathways encouraged us to ask if these features can be used to associate a mutated gene to each compound affecting this gene’s pathway. For this purpose, we trained a multilayer perceptron (MLP) with a single hidden layer (**Materials and Methods**) using the DL-derived feature vectors as input and the gene corresponding to the disrupted pathway as label, based on images of drug-treated bacteria across the three replicates (**Fig. 4d**). Images of bacteria exposed to the four DNA gyrase inhibitors BDM71403, ciprofloxacin, gepotidacin and moxifloxacin were correctly associated to either *gyrA* or *gyrB* depletion mutants. The two PBP inhibitors ampicillin and carbenicillin were in turn associated to *sepF,* ∆*steAB* or *∆glpR*, mutants which are known to perturb the division and thus the septal cell wall machineries. Ethambutol was correctly associated to the cell elongation defective mutant *wag31*.

In summary, these results show that our DL model links mutants to drugs that perturb similar pathways directly from high-throughput images. The possibility of identifying the biological pathway disrupted by a drug from images alone represents an important step towards mutant-based target deconvolution of hit compounds by comparing drug-induced phenotypes to those found in genome-wide mutant libraries.

### Detection of mechanistic fingerprints in cell cycle associated pathways

Based on the results above, wherein our DL model successfully linked phenotypes of drug-treated bacteria to mutants of the targeted pathways, we asked if a similar approach could be used to provide insights into fundamental biological processes. As a proof of principle, we focused on five of the above-mentioned mutants implicated in the bacterial cell cycle and for which functional data is available in the literature (**Fig. 5a**). These included the three knockout strains ∆*steAB*, ∆*glp* and ∆*glpR* and the two conditional depletion mutants *sepF* and *wag31*, which were imaged at four time points (3, 6, 9 and 24h) corresponding to increasing depletion levels. During Wag31 depletion, cellular morphologies evolved from rod to ovoid and finally to coccoid shape^38^ (**Fig. 5b**). Upon SepF depletion, the Z-ring disassembles, and cells start to elongate from roughly 2μm in WT to over 10μm during depletion^35^. These elongated cells eventually branch (**Fig. 5b**) – which corresponds to uncontrolled and aberrant pole formation – and finally lyse. Glp and GlpR form a tight complex at the septum and respectively bind FtsZ and Wag31, thus linking the division and elongation cytoskeleta at the septum to ensure the correct transition between division and elongation. Glp depletion results in elongated cells, while GlpR depletion resembles WT cells in classical morphological readouts^37^ (**Fig 5b**). Lastly, SteAB forms a complex involved in cell wall remodelling by regulating PG hydrolysis and is also responsible for correct onset of cytokinesis^36,40^. ∆*steAB* cells are elongated and usually display multiple septa^36,41^ (**Fig 5b**).

We imaged the above-mentioned mutants in triplicates as previously described and used our DL model trained on antibiotics (**Table 1**) to obtain feature vectors which showed low plate-to-plate variation across replicates (**Supp Fig. 7b**). Plotting the extracted feature vectors with UMAP revealed clustering of mutants consistent with different stages of the bacterial cell cycle (**Fig. 5c**). Early-time-point mutants (3, 6 and 9h) of Wag31 formed a tight cluster and were the closest to the WT cells compared to all other mutants. This is consistent with the fact that early depletion of *wag31* leads to a relatively subtle but significant phenotype of cellular rounding^38^. The late time-point mutant of *wag31* (24h) localised away from all other conditions reflecting its distinctly rounded shape. All *sepF* mutants, except for the earliest time-point at 3h, formed a cluster that was the furthest away from WT cells, reflecting the extreme elongation phenotype induced by interfering with FtsZ function^35^. The early-time-point sepF mutant was near the three knockout mutants ∆*steAB*, ∆*glp* and ∆*glpR*, known to be perturbed in their division machinery. Interestingly, the ∆*glpR* mutant, which did not differ significantly from WT cells when analysed using segmentation and classical morphological features^37^, clustered away from WT and together with the ∆*glp* mutant. Glp and GlpR are known partners *in vivo* where they localise to the septum during cell division. They have also been shown to form a strong complex *in vitro* (nM range affinities)^37^. This suggests that the DL model can pick up relevant physiological signatures overlooked by classical cell analytical methods.

To further quantify these observations, we constructed an adjacency graph (**Fig. 5d**) from a cosine similarity matrix computed on median feature vectors (**Supp. Fig. 7c**) setting the threshold such that each node was linked by at least one edge (**Materials and Methods**). Mutants disrupting cell elongation and cell division occupied the extremes with higher depletion levels being the furthest away from all other conditions. Mutants associated with the division machinery were localised between these two extremes. Notably, our adjacency graph is in quite good agreement with the known function of these proteins in the bacterial cell cycle, as shown in **Fig. 5a**. For example, Δ*steAB* and Δ*glp* both lead to multiseptated elongated phenotypes that prevent correct cell cycle progression^36,37^ and Δ*glpR* falls in between Δ*glp* and WT.

Taken together, these results show that feature vectors obtained with our DL model from images of mutant strains recover known mechanistic relationships between bacterial genes thus paving the way towards genome-wide studies of bacterial gene function directly from imaging data.

## Discussion

In this work, we present a DL-based analysis approach to extract highly specific features from high-throughput images of chemically and genetically perturbed *Cglu* bacteria. We show that these features enable robust discrimination between drugs by MoA, recognition of the MoA of previously unseen drugs and detection of drugs with a novel MoA. In addition, we find that our DL model correctly classifies all but one drug at a concentration above 5xMIC. Then, we demonstrate that the learned features allow to link pathway-specific chemical perturbations to genetic perturbations of the same pathway for three pathways commonly targeted by antibacterial drugs: DNA gyrase, elongation and division/cell wall machineries. Finally, we show encouraging results towards annotating bacterial gene function directly from images using the cell cycle of *Cglu* as a case study. In particular, our approach recovers a known relationship between genes (*glp* and *glpR*) while this was not possible with traditional image-based profiling^37^.

We used *Cglu* as a surrogate model to develop a pipeline to screen for TB inhibitors. Ideally, screens should be carried out using the pathogen in question. However, in this regard not all bacteria are equal. While traditional bacterial cytological profiling (BCP) is sufficient to determine the MoA in *E. coli* or *Bacillus subtilis*, this is not the case for *Mtb*^13,14,16,17^. BCP performs poorly on mycobacteria^16,17^ and DL methods have only recently been employed to successfully distinguish drug-induced phenotypes in *Mtb*^17^. A key obstacle of using classical methods to profile *Mtb* is the intrinsic cellular heterogeneity of the tubercle bacillus. Indeed, elegant work by Smith *et al*. demonstrated that standard phenotypic analysis is not sufficient to distinguish drug-induced phenotypes of *Mycobacteria*^16^. To address this, the authors developed a complex pipeline that takes into account cellular heterogeneity and relies on iterative feature selection including variables linked to cell-to-cell variation. Although their pipeline can address detailed questions around known MoAs and off-target effects of a small number of compounds, it would be challenging for high-throughput screening of large compound libraries. Moreover, while these methods enable MoA recognition of hit compounds with respect to known compounds, they are unable to identify drug targets of compounds that exhibit novel MoAs. Here, we show that we can identify novel/unseen MoAs and we address the above limitation by linking drugs to mutants where similar pathways have been disrupted directly from high-throughput images (**Fig. 4**).

Another potential limitation of these prior studies is the choice of model organism. To overcome the limitations of working on *Mtb* directly while keeping a parent model that retains core conserved cellular pathways, we chose the industrially relevant bacterium *Cglu*. It is not pathogenic, significantly simplifying the experimental pipeline and grows rapidly (doubling time 1h) in defined media. This is a crucial consideration when producing data for DL-based analysis where reproducible phenotypic readouts and minimal batch-to-batch variation are key, as even slight variations in media composition can lead to important morphological alterations^42^. Additionally, drug-induced phenotypes are more homogenous when compared to *Mtb* thereby simplifying the analysis. We therefore believe that *Cglu* is an attractive model organism to screen for novel TB drugs. In particular, the similar and unique cell wall composition (a major target for anti-TB treatment^43^) and the conserved core machinery across the two species are important advantages. We do however acknowledge that relevant TB targets in this surrogate model will be limited to conserved mechanisms and is thus not suited for finding *Mtb-*specific pathways such as those linked, for example, to virulence.

In prior work, we showed that brightfield images alone are sufficient to distinguish between drug MoAs in *E. coli* bacteria^31^. Despite the absence of clearly discernible visual phenotypes in most conditions (**Fig. 2a**), our DL model produced good results on the dataset of *Cglu* bacteria only using the brightfield channel (**Supp. Fig. 8a**). However, the addition of fluorescent dyes improved prediction accuracies (79.16±7.25% vs. 86.78±7.90% computed per FOV; **Supp. Fig. 8b**). This is likely due to drug-treated *Cglu* bacteria exhibiting much subtler phenotypes compared to *E. coli*.

In the context of a phenotypic screen, early identification of the hit’s MoA is crucial to select promising compounds for further downstream development. Since our approach can recognise the MoA of previously unseen compounds directly from images by comparison to drugs with known MoA or detect that they have a novel MoA (**Fig. 3**), it sidesteps laborious experiments in the lab. By providing insights on the MoA directly from imaging data generated in a screen, our method is a step towards rational hit selection. Our data also suggest that compounds with previously undiscovered MoAs could quickly be detected and their target pathways identified by comparing their phenotypes to those of mutants.

We also observed that predictions from our DL model were correct for all but one drug above 5xMIC and for a subset of drugs MoA predictions were correct regardless of concentration (**Supp. Fig. 4a**). Notably, if these drugs had been used in the context of a screen at 10μM, the MoA of only two (clofazimine and ethambutol) out of the 14 active drugs with a previously seen MoA would not have been correctly identified. While this is encouraging, it should be noted that these drugs have undergone extensive lead optimisation and are thus more potent than typical hit compounds. It therefore remains to be explored how DL performs on large non-specific compound libraries.

Another direct application of this work is the possibility for an immediate validation of compound series resulting from medicinal chemistry. Compounds can be rapidly tested for both cell penetrability and efficacy on the target, as well as possible off-target effects resulting from chemical modifications. Since our DL model encodes dose-dependent phenotypic changes of drug-treated bacteria on MoA-specific trajectories in the feature space (**Supp. Fig. 4b**), we speculate that our approach might also be used to aid lead optimisation of hit compounds through image-based structure activity relationship (SAR) studies. SAR landscapes could be constructed based on the features extracted by our DL model from images of drug-treated bacteria to map the potency of a given change in molecular structure onto these dose-dependent trajectories. This could enable high-throughput studies where the molecular structure of a lead compound is systematically altered to gain insights on potency directly from images.

Making a direct link between mutants and drugs is only possible in very few well documented cases such as DNA gyrase inhibitors for which the target is possibly unique and experimental structures of drug-target complexes are available^44^. In contrast, in the case of PBPs one drug may target several enzymes, and mutant strains will not reflect a one-target-one-drug scenario. Other drugs used in the clinic have controversial target attributions and many MoAs cover global physiological perturbations, as exemplified by clofazimine, which at least in part targets the respiratory chain, but has also been reported to have more general effects on disrupting membrane potential and homeostasis^29^. Dissecting these possible effects will be of great importance, not only for MoA detection but also for quantifying off-target effects that could be more general and lead to toxicity. While our results (**Fig. 4**) remain to be validated on larger datasets, we believe that they point towards a strategy of target deconvolution by relating drug-induced phenotypes to whole-genome mutant libraries.

Our results have also shown that we can not only detect the dramatically different phenotypes linked to a complete disruption of cell division (SepF) or elongation (Wag31), but also more subtle phenotypes that cannot be distinguished in classical segmentation-based feature analysis (**Fig. 5**). One such example is the GlpR mutant strain that has a WT morphology when segmented and analysed for cell parameters such as length and width^37^. Our analysis, however, reveals clear differences to the WT condition (**Fig. 5c**) and high similarity to the Glp strain, in line with the known biological function of GlpR. Indeed, GlpR has been shown to form a strong complex with Glp (nM range) at the corynebacterial septum during cell division^37^. This shows that our DL model picks up functionally relevant features that escaped classical analysis.

In conclusion, we have demonstrated that images of bacteria contain a high degree of pathway-specific phenotypic features and have introduced a DL approach capable of extracting this rich information source. With growing datasets facilitated by automated screening protocols, methods like the one presented herein are poised to greatly speed up the discovery of novel antibiotics and chart a path towards systematically investigating bacterial gene function directly from imaging data.

## Materials and Methods

### Bacterial strains, culture conditions and antibiotic treatment

*C. glutamicum* ATCC 13032 (*Cglu*) was used as a wild-type (WT) strain. *Cglu* was grown overnight in a flask in CGXII medium^45^ with 4% sucrose at 30°C and 110 rpm shaking. The following day, the culture was diluted to OD_600_ of 1.3 in a flask in CGXII medium with 4% sucrose, grown for 2.5 h at 30°C and 110 rpm (to a required OD_600_ of 2), and then dispatched in a 96-well plate containing randomised antibiotics. Cells were then treated for 16 h at 30°C and 110 rpm, before being fixed as described below. *Cglu_P_ino_-wag31* and *Cglu_P_ino_-sepF* strains were inoculated in CGXII 4% sucrose and grown for 8 h at 30°C 110rpm, then diluted to OD_600_ of 1.3 in CGXII 4% sucrose supplemented with 1% myo-inositol and grown for either 3, 6, 9 or 18 h at 30°C 110rpm for different depletion levels. *Cglu_P_ino_-GA* and *Cglu_P_ino_-GB* were inoculated in CGXII 4% sucrose and grown overnight at 30°C 110rpm, diluted the next day to OD_600_ of 1.3 in CGXII 4% sucrose 1% myo-inositol, and grown at 30°C 110rpm for 7 h 30 min. Cultures were then diluted again to an OD600 of 1.3 in fresh CGXII medium with 4% sucrose 1% myo-inositol and grown for 18 h at 30°C 110rpm to reach total depletion. *Cglu_Δglp*, *Cglu_ΔglpR* and *Cglu_ΔsteAB* were inoculated in CGXII 4% sucrose and grown overnight at 30°C 110rpm, diluted the next day to OD_600_ of 1.3 in CGXII 4% sucrose and grown for 5 h at 30°C 110rpm.

### Antibiotics dispatching

For the dataset used for training, antibiotics were dispensed by acoustic droplet ejection using an Echo 550 liquid handler (Beckman Coulter Life Sciences) into dry 96-well plates (clear, flat bottom, TPP 92097). Well positions for each assay plate were randomised prior to the Echo run; randomised pick lists were generated upstream and imported into Echo Cherry Pick to create the transfer instruction files. Compounds from the FDA-approved & Passed Phase I Drug Library (Selleckchem L3800) included amoxicillin, ampicillin, cefotaxim, doxycycline, moxifloxacin, rifabutin, rifampicin, linezolid, carbenicillin, clarithromycin, trimethoprim, clofazimine, ethambutol, sulfamethoxazole, ciprofloxacin, amikacin, kanamycin, streptomycin. Additional compounds were gepotidacin (MCE HY-16742) novobiocin (Sigma N1628) and BDM71403^22^. All antibiotics were dispensed from DMSO stocks except novobiocin, amikacin, kanamycin, and streptomycin, which were dispensed from aqueous stocks. For each antibiotic, four concentrations corresponding to 0.5, 1, 5, and 10xMIC were dispensed. Wells containing DMSO were backfilled to achieve a uniform final DMSO content of 0.49% in the assay. Control wells included six vehicle controls with 0.49% DMSO and six vehicle controls with water. Immediately after acoustic transfer, 10 µL of growth medium were added to each well to minimize evaporation and compound degradation, followed by culture addition.

For the manual dataset imaged together with the mutant strains (**Fig. 4**), all dilutions were made through manual pipetting. Moxifloxacin (SIGMA SML1581), BDM71403^22^, gepotidacin (MCE HY-16742) and rifampicin (Sigma R3501) were diluted in DMSO and added to the culture at a final DMSO concentration of 0.1%. Ciprofloxacin (SIGMA PHR1044), ethambutol (Sigma 1070-11-7), carbenicillin (Euromedex 1039A), ampicillin (Sigma A9518), amikacin (PHR1654) and kanamycin (K0254-20ML) were diluted in water. Eight control wells were filled with 0.1% of DMSO and eight control wells with water. Concentrations were used as 0.5, 1, 5 and 10xMIC values listed in **Table 1**.

### Minimum inhibitory concentration determination

The MIC determination against wild-type *Cglu* ATCC13032 were determined by agar dilution method on MH agar and broth. For the agar dilution method, MIC values were obtained using the previously described proportion method^46^. Briefly, 10^3^ and 10^5^ CFU were inoculated onto MH agar containing two-fold serial dilutions of each compound. After 1 day of incubation at 30°C, colonies were enumerated. The MIC was defined as the lowest concentration of antibiotic resulting in complete inhibition of growth or in growth of fewer than 1% of the inoculum. For the broth dilution method, antibiotics were prepared in two-fold serial dilutions in a 96-wells flat-bottom plate, each well containing 100 µL of MH broth with the corresponding antibiotic. Wells were inoculated with 2 µL of an overnight culture. Plates were incubated for 24h at 30°C with 600 rpm shaking. OD_600_ values were measured at 24h. MIC was determined as absence of growth compared to blank.

### Sample fixation and preparation for imaging

For 96-well plates, 75 µL of bacterial cultures were mixed with 180 µL of ice cold EtOH and stored for 1 h at -20°C. Samples were washed 3 times with 500 µL, 700 µL and 75 µL of filtered sterile PBS, and diluted to OD_600_ of 0.025 in PBS supplemented with FM4-64 (Invitrogen T13320) 1ug/mL final and Hoechst 33342 (cell signaling technology #4082) 1 µg/mL final. 100 µL of each sample were loaded in a PhenoPlate (Revvity 6055302), which were stored at 4°C and centrifuged before imaging. Those steps were either done manually either using an Apricot PP5 robot (sptlabtech). For mutant samples, 300 µL of each culture were harvested and fixed with 700 µL of EtOH ice cold for 1 h at -20°C. Samples were then washed 3 times in 300 µL of filtered sterile PBS, and diluted to OD_600_ of 0.025 in PBS supplemented with FM4-64 (Invitrogen T13320) 1 µg/mL final and Hoechst 33342 (cell signaling technology #4082) 1 µg/mL final. 100 µL of each sample were loaded in a PhenoPlate (Revvity 6055302) following a random map. Plates were stored at 4°C and centrifuged before imaging.

### High-throughput imaging

Plates that were stored at 4°C were left at least 30 min at room temperature and centrifuged 10 min at 500rpm. An Opera Phenix Plus (Revvity) was used in widefield mode to image 50 fields of view per well, with a 63/1.15 water-immersion objective. The fluorescent dyes – Hoechst 33342 and FM4-64 – were acquired simultaneously with a 405 nm or 488 nm excitation laser lines and a 435-480 nm or 650-680 nm bandpass filters, respectively, in addition to brightfield imaging.

### Image preprocessing

Images were converted from 16-bit to 8-bit and pixel intensities clipped by percentile between 0.1 and 99.9 to remove spurious saturated pixels. Images from five out of six plates were randomly split into training (80%) and validation (20%) set. Data augmentation was applied to full-resolution images (2,160x2,160 pixels) on the fly (random flipping and rotation, random changes in brightness, contrast, saturation and hue). Randomly rotated images were then centre-cropped (1,528x1,528 pixels) to account for potential edge effects. Next, a random crop (1,500x1,500 pixels) was obtained and divided into 9 tiles (500x500 pixels) which were resized to 256x256 pixels and pixel intensities normalised between -1 and 1.

### CNN architecture and training

A custom CNN was implemented in PyTorch v2.3.1 based on an EfficientNet-B0^47^ backbone. The CNN was first pre-trained for 100 epochs using an Adam^48^ optimiser with a learning rate of 0.001 and an effective batch size of 144 images (16 images divided into 9 tiles). Image tiles were passed through the backbone, with weights initialised by pre-training on ImageNet, and latent embedding vectors corresponding to individual image tiles aggregated by averaging to obtain a global embedding vector for each input image of size 1,500x1,500 pixels. For pre-training, the model was trained to predict the combination of drugs and concentrations as a multi-class classification task using a weighted cross-entropy loss (classes were inversely weighted by the number of images in the training data).

For fine-tuning, model weights corresponding to the lowest validation loss during pretraining were chosen and only images corresponding to drug treatments at the highest concentration (10xMIC) were used for training. Images of bacteria exposed to drugs belonging to the same MoA were grouped into the same class. During fine-tuning, models were trained using a combination of three loss terms with latent embedding vectors obtained from an intermediate fully connected layer of 16 neurons. First, a triplet loss^49^ was used to explicitly promote the formation of clusters by MoA in latent space, defined as:

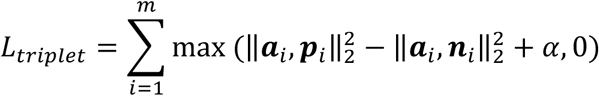

where 𝒂_i_ denotes the 𝑖th anchor embedding vector, 𝒑_i_ denotes the 𝑖th embedding vector of a positive example and 𝒏_i_ denotes the 𝑖th embedding vector of a negative example, while 𝑚 corresponds to the number of hard triplets. The margin 𝛼 was set to 0.1. During training, triplets were assembled on the fly based on their MoA class. In addition, hard triplet mining was applied, such that the loss was only computed on triplets where the negative example was closer to the anchor than the positive example. Therefore, hard triplets satisfy the following inequality:

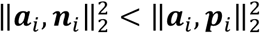

Second, a centre loss^50^ was used to increase the tightness of clusters. The centre loss seeks to minimise intra-class variations while ensuring that different classes remain separable and is thus defined as:

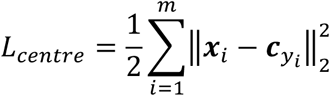

where 𝒙_i_ denotes the 𝑖th embedding vector, 𝒄*_yi_* denotes the centre of the 𝑦_#_th class corresponding to the MoA of a given embedding and 𝑚 is the effective batch size.

Third, a weighted cross-entropy loss was used to obtain MoA predictions (weighted by the inverse number of training images per class). The final loss function used for training was thus:

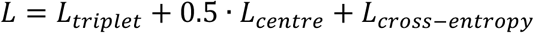

### Hierarchical clustering

Hierarchical clustering was performed on 16-dimensional feature vectors with Scipy v1.12.0 using the ‘scipy.cluster.hierarchy’ function. The distance between clusters was computed with the Ward variance minimization algorithm (‘ward’). The clustering threshold was set with respect to the negative control conditions (Water and DMSO).

Hierarchical clustering on cosine similarity matrices was obtained with the ‘seaborn.clustermap’ function from Seaborn v0.11.2. using the ‘scipy.cluster.hierarchy’ function computed on 16-dimensional feature vectors. The distance between clusters was computed with the Farthest Point Algorithm (‘complete’) and a cosine distance metric.

### Local outlier factor algorithm

Embedding vectors from images of previously seen conditions were obtained from the hold-out test plate. A local outlier factor classifier was applied to these embeddings using all available concentrations with the scikit-learn package in Python 3.9. The number of k neighbours was set to k=10 and Euclidean distance was used as the distance metric. Outlier scores were computed by taking the fraction of detected outliers considering all fields of view (FoVs) for a given condition.

### Multilayer perceptron

Embedding vectors of 16-dimensions were obtained from images of mutants across three replicates. The ‘sklearn.neural_network.MLPClassifier’ function from scikit-learn v1.5.1 was then used to construct an MLP with a single hidden layer of 64 neurons using a ReLU non-linearity. The MLP was trained for 200 iterations with Adam and a learning rate of 0.001. Predictions were then obtained from 16-dimensional embedding vectors of images of drug-treated bacteria.

### Adjacency graph

A cosine similarity matrix was first computed on 16-dimensional median embedding vectors. Then, a binary mask was obtained by thresholding the cosine similarity matrix. The threshold was set such that each node was connected by at least one edge. The element-wise product between the mask and the cosine similarity matrix was then computed to obtain a masked cosine similarity matrix. To obtain an adjacency matrix, the diagonal of this masked cosine similarity matrix was set to 0. An adjacency graph was then constructed from the adjacency matrix with the networkx v3.2.1 package using a spring layout.

## Acknowledgements

We gratefully acknowledge the C2RT core facilities at the Institut Pasteur, in particular UtechS-PBI and PF-CCB, and funding from Institut Pasteur ATC-DDS (Technological Targeted Actions – Drug Discovery & Screening) for compound library purchase. The UTechS Photonic BioImaging, C2RT, Institut Pasteur, is supported by the French National Research Agency (France BioImaging, ANR-24-INBS-0005 FBI (BIOGEN); Investments for the Future) and acknowledges Institut Pasteur and the Région Île-de-France (DIM1Health program) funding for the use of the Opera Phenix system. This work was supported in part by grants from the Agence Nationale de la Recherche (ANR, France), contracts ANR-21-CE11-0003 (A.M.W.), ANR-24-CE11-4058 (A.M.W), Fondation pour la Recherche Médicale, FRM, EQU202303016284 (P.M.A.), Institut Pasteur PTR_726_BactImMorph (A.M.W.) and by institutional grants from the Institut Pasteur, the CNRS, and Université Paris Cité. J.P. was partially funded through the AMX program from the École Polytechnique. D.K. was funded by the Pasteur-Paris University International doctoral program, the INCEPTION program (Investissement d’Avenir grant ANR-16-CONV-0005) and a Fondation pour la Recherche Médicale Fin de Thèse grant (FDT202404018132). We also acknowledge the INCEPTION program for funding a GPU farm used in this work. We thank Cédric Thépenier, Kelvin Kho, Ivo Gomperts Boneca and Spencer Shorte for helpful discussions.

## Author contributions

DK, JP, PMA, SP, CZ, and AMW designed the research. JP designed and established the protocols and prepared all the samples for high content analysis. YMB carried out additional microbiology experiments. NM and NA provided advice on the experimental settings for high content imaging and carried out the acquisitions. ES and AA performed MIC determinations. SP provided expertise and reagents for the DNA gyrase experiments. DK designed and carried out all the computational analysis. JC and AZ oversaw the automated compound delivery. DK, JP and AMW wrote the paper. All authors edited the paper.

