## Supplementary Material for "Deep learning extracts MoA-specific signatures from high-throughput images of chemically and genetically perturbed *Corynebacteria*"

*These authors contributed equally

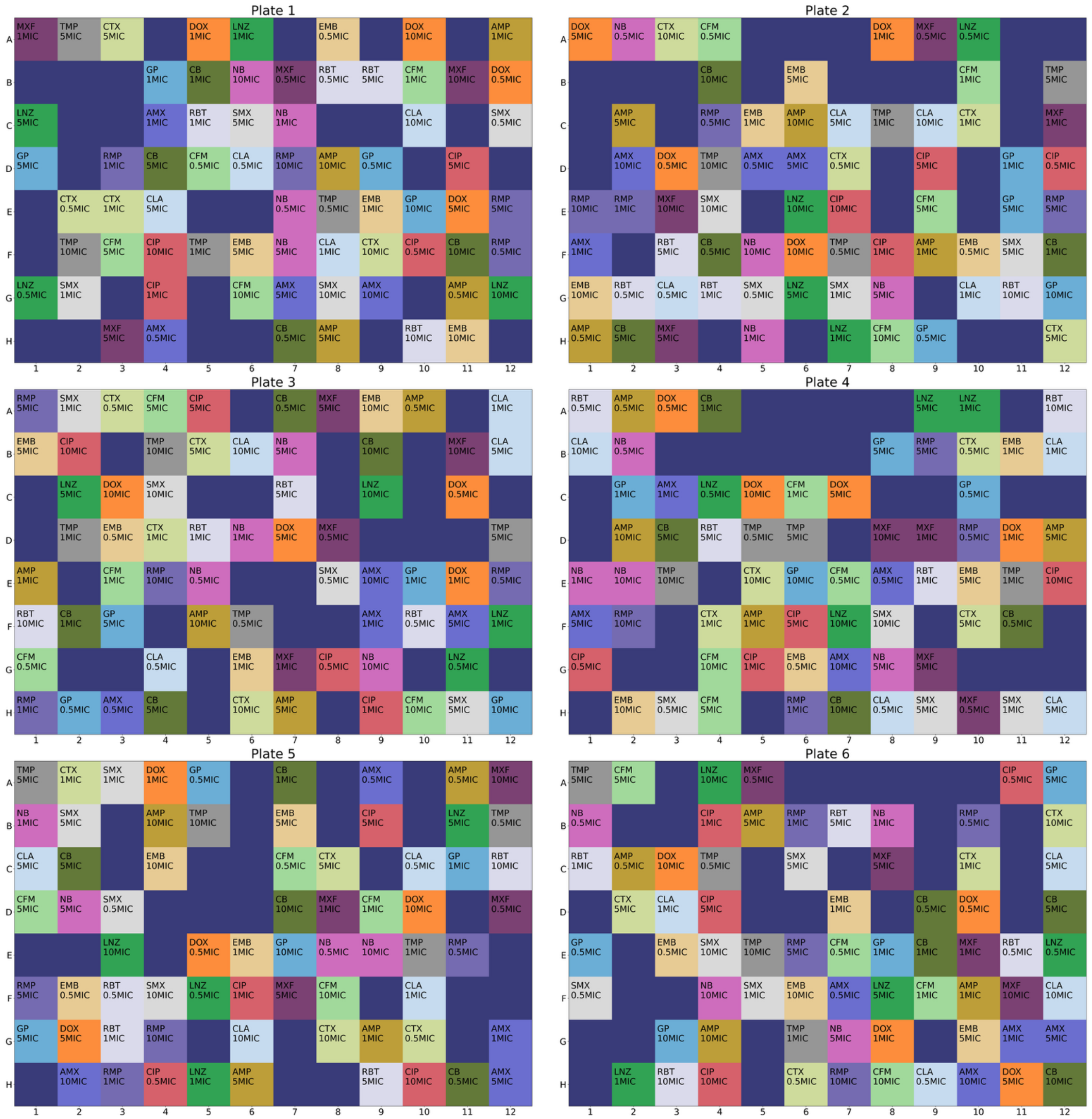


**Supp. Fig. 1. Plate layouts.** Annotated plate layouts with drug concentration shown below. A total of 17 drugs at four different concentrations were distributed with an acoustic liquid handler. Amoxicillin: AMX, carbenicillin: CB, cefotaxime: CTX, ampicillin: AMP, ethambutol: EMB, ciprofloxacin: CIP, moxifloxacin: MXF, novobiocin: NB, gepotidacin: GP, clarithromycin: CLA, doxycycline: DOX, linezolid: LNZ, clofazimine: CFM, rifampicin: RMP, rifabutin: RBT, trimethoprim: TMP, sulfamethoxazole: SMX. Empty positions represent controls (water and DMSO).

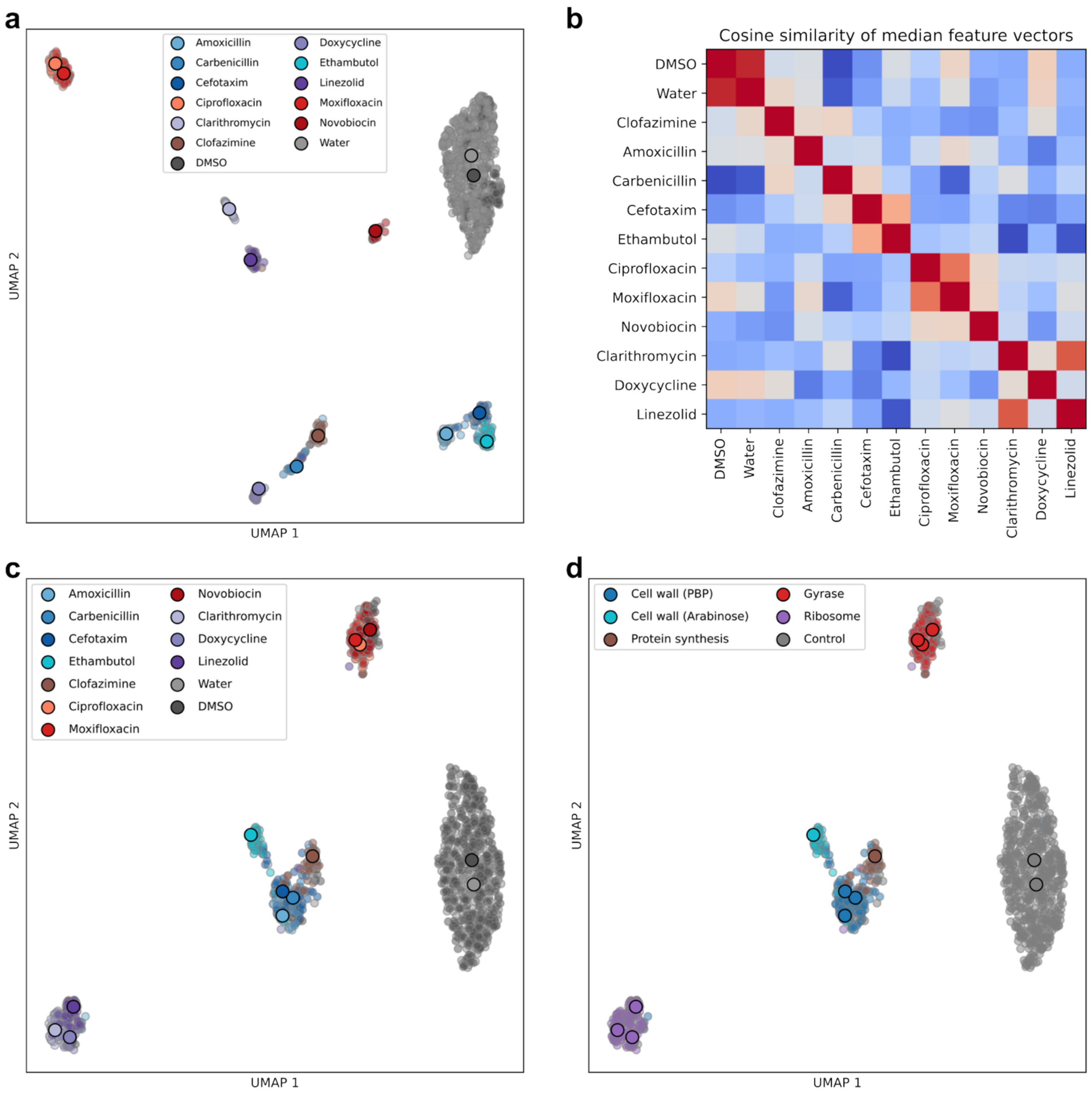


**Supp. Fig. 2. Comparison of feature vectors before and after fine-tuning.** (**a**) UMAP plot of feature vectors obtained from images of drug-treated bacteria at 10xMIC from a model trained on the pretext task of classifying the combination of antibiotic and concentration without further fine-tuning (**Materials and Methods∫)**. (**b**) The cosine similarity matrix was computed on median feature vectors that were obtained from images of drug-treated bacteria using a model that did not undergo fine-tuning (**Materials and Methods**). (**c, d**) UMAP plot of feature vectors using a model that was further fine-tuned to predict the MoA of antibiotics at 10xMIC coloured by drug label (**c**) and MoA (**d**). Small dots correspond to individual FOVs and large dots correspond to the median feature vector of a given condition.

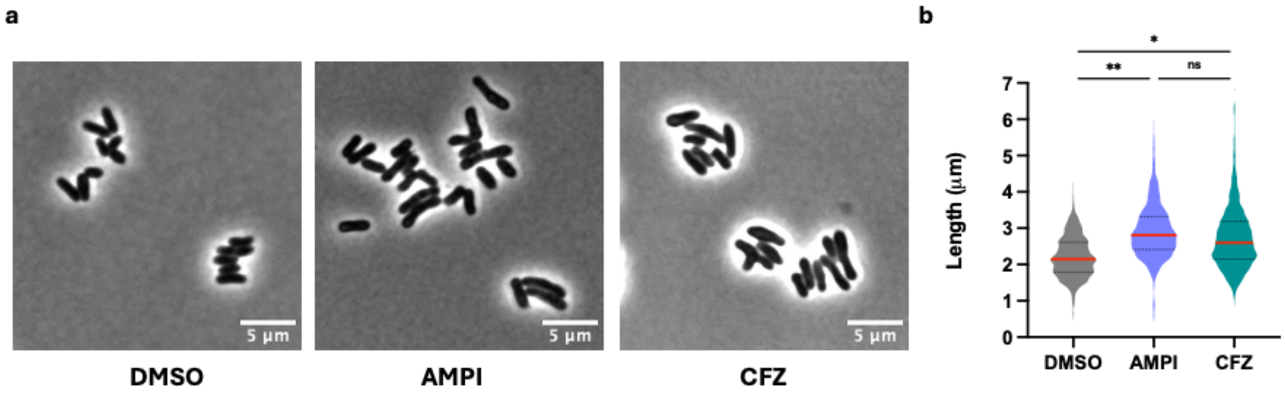


**Supp. Fig. 3. Comparison of *Cglu* morphologies exposed to clofazimine and ampicillin.** *Cglu* cultures were inoculated at $OD_{600}$=1. After 2 h of growth, antibiotics were added and cultures and were incubated for 18 h. Bacteria were treated with antibiotics following same protocol as the one used to generate the high-throughput image datasets (**Materials and Methods**). (**a**) Phase contrast images were acquired for 100 ms. Scale bars, 5 µm. (**b**) Violin plots show the distribution of cell length (between bacterial cells treated with DMSO (n = 1519), ampicillin (AMPI; n = 1100) and clofazimine (CFZ; n = 1253). Cohen's d: DMSO/AMP, **d = 1.05; DMSO/CFZ, *d = 0.72; AMPI/CFZ, ns = 0.2.

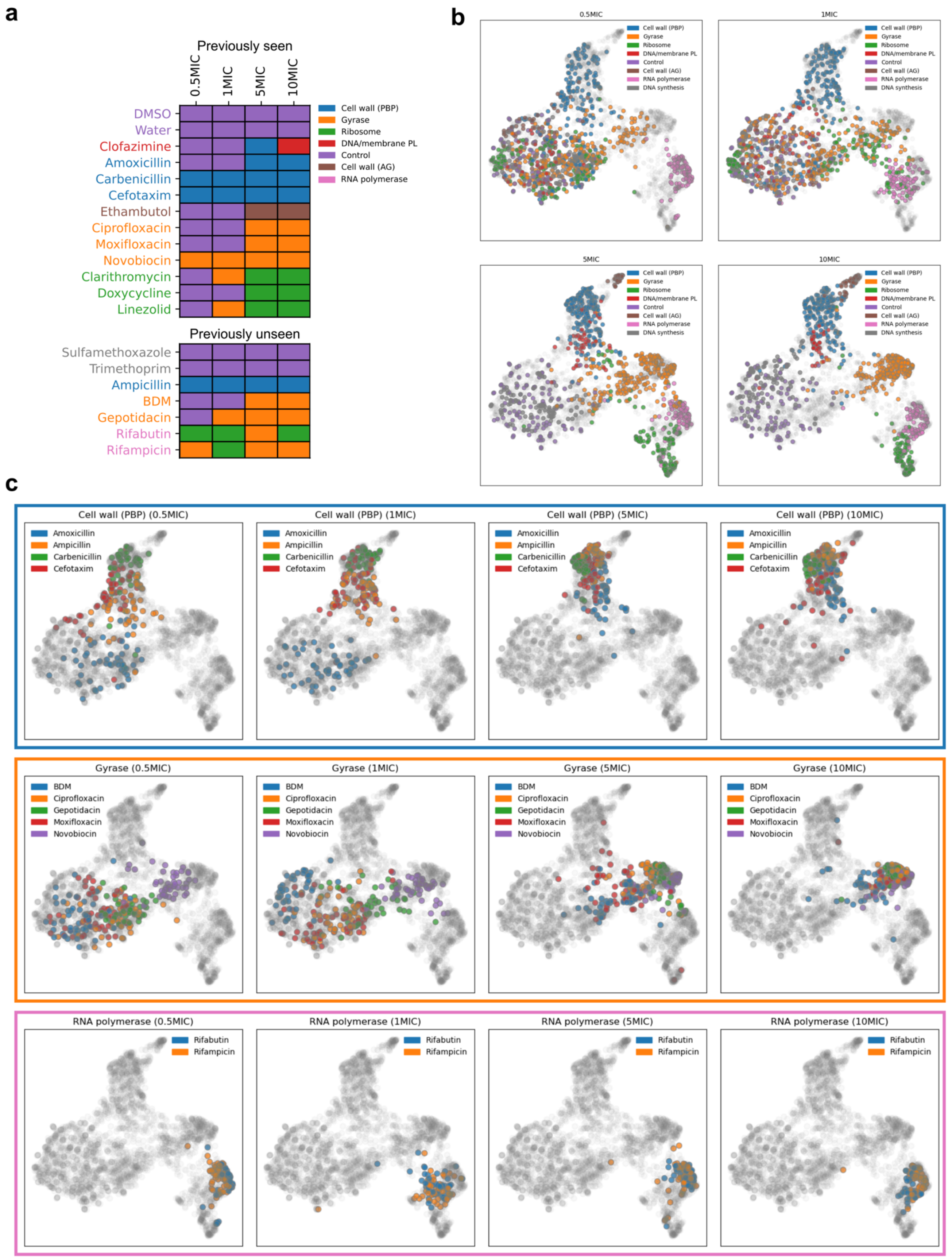


**Supp. Fig. 4. Concentration-dependent effects.** (**a**) Well-level MoA predictions obtained from images of bacteria exposed to both previously seen and unseen antibiotics at four different concentrations on a hold-out test plate with a DL model fine-tuned on images of bacteria exposed to antibiotics at 10xMIC. (**b, c**) Lower-dimensional projection of feature vectors from images of bacteria exposed to antibiotics at four concentrations was obtained with UMAP. Dots correspond to individual images and colours to the MoA of a given antibiotic. Dose-dependent latent trajectories can be observed with feature vectors tending to localise further away from control conditions with increasing concentration.

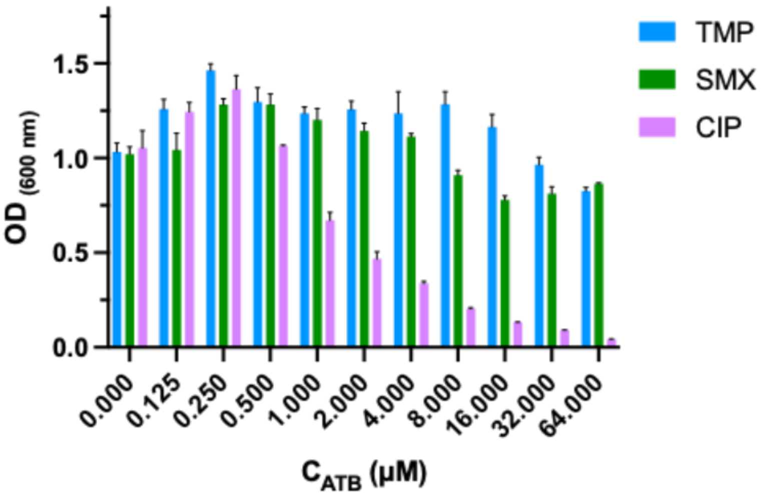


**Supp. Fig. 5. Trimethoprim and Sulfamethoxazole are inactive in *Cglu.*** *Cglu* was inoculated at a final $OD_{600}$= 0.2 in a 96-well plate and exposed to antibiotics (trimethoprim TMP, sulfamethoxazole SMX and ciprofloxacin CIP) at different concentrations (as indicated). Bacterial growth was recorded after 18 h and quantified by $OD_{600}$ measurements. Bars indicate mean values with error bars corresponding to mean ± sd.

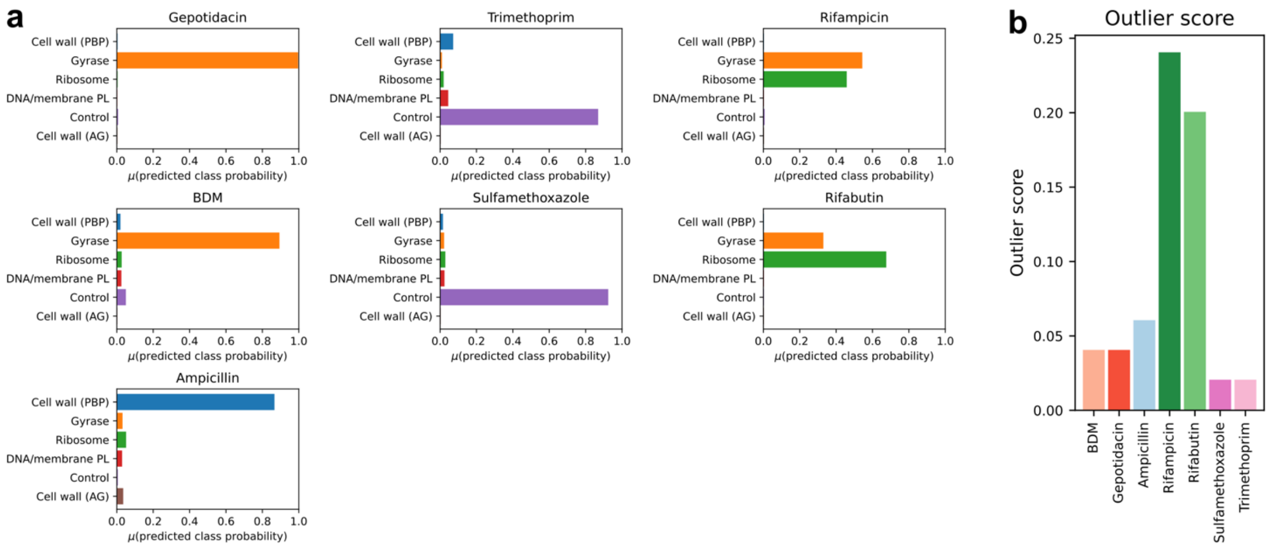


**Supp. Fig. 6. Predictions on previously unseen drugs and outlier scores.** (**a**) Mean classification probabilities averaged across all fields of view (FOVs) are shown for each of the previously unseen drugs. (**b**) Outlier scores for previously unseen drugs obtained with a local outlier factor algorithm (**Materials and Mehods**).

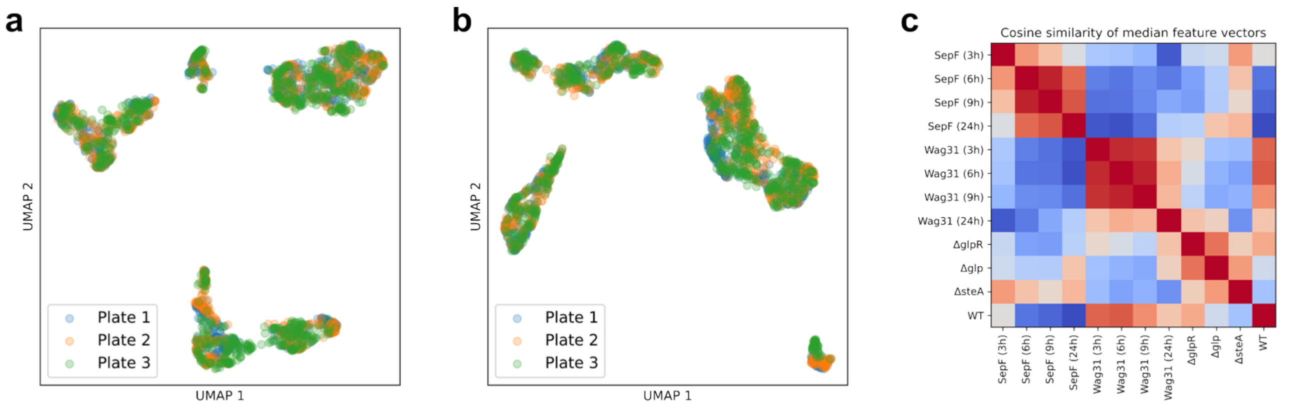


**Supp. Fig. 7. Model performance.** (**a**,**b**) UMAP of feature vectors coloured by plate showing good agreement across replicates for data shown in (**a**) **Figure 4b** and (**b**) **Figure 5c**. (**c**) Cosine similarity matrix computed on median feature vectors of mutants from which the adjacency matrix in **Figure 5d** is obtained.

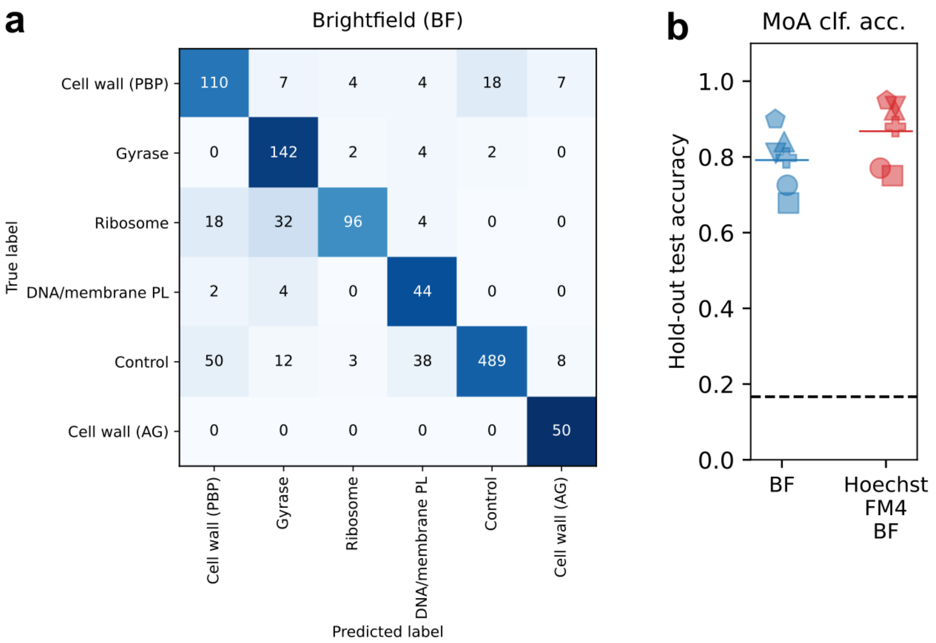


**Supp. Fig. 8. Model performance for different choices of imaging channels.** (**a**) Confusion matrix on a hold-out test plate for a model only trained using the brightfield channel at 10xMIC. Values within the matrix indicate the number of FOVs. (**b**) Cross-validated MoA classification performance for models trained only using brightfield (BF, blue) and brightfield in combination with Hoechst and FM4-64 (Hoechst FM4 BF, red) at 10xMIC.

**Supp. Table 1:** Bacterial mutant strains

| **Strain** | **Genetic modification** | **Affected gene** | **Affected pathway** |
|---|---|---|---|
| *Cglu_WT* | ∅ | ∅ | ∅ wild-type rod-shaped |
| *Cglu_*$P_{ino}$*-wag31* | Conditional depletion | cg2361 | Elongation defect (no elongation, ovoid to coccoid) |
| *Cglu_*$P_{ino}$*-sepF* | Conditional depletion | cg2363 | Early division defect (no division, elongated to branched) |
| *Cglu_*$P_{ino}$*-GA* | Conditional depletion of *gyrA* | cg0015 | DNA segregation defect, diffuse nucleoid |
| *Cglu_*$P_{ino}$*-GB* | Conditional depletion of *gyrB* | cg0007 | DNA segregation defect, diffuse nucleoid |
| *Cglu_Δglp* | Knockout | cg1005 | Division defect (multiseptal, elongated) |
| *Cglu_ΔglpR* | Knockout | cg1003 | Wild-type-like rod-shaped |
| *Cglu_ΔsteAB* | Knockout | cg1603/ cg1604 | Cell wall defect (multiseptal, elongated) |
